# Initial sample processing can influence the soil microbial metabarcoding surveys, revealed by *Leucocalocybe mongolica* fairy ring ecosystem

**DOI:** 10.1101/2020.06.09.142018

**Authors:** Mingzheng Duan, Tolgor Bau

## Abstract

In this study, we aimed to investigate the influence of soil preservation approaches, especially cryopreservation and high temperature-drying on the sequencing quality of its microbial community and the background microbial diversity information of fairy ring soil from *Leucocalocybe mongolica*. Through DNA metabarcoding surveys based on 16S rDNA and ITS barcodes, we observed that the bacterial abundance was notably changed when the soil samples were exposed in room temperature for 4 hours, whereas the fungal composition was not significantly changed. Moreover, the soil samples preserved their major microbial structures even after high temperature-drying for 12 hours, whereas their microbial diversity was influenced. Overall, a total of 9283 and 1871 OTUs were obtained from soil bacteria and fungi, respectively, and we observed that Chthoniobacteraceae and Tricholomataceae were the dominant bacterial and fungal families in the fairy ring soil, respectively. Our study reveals the impact of soil processing methods on the microbial community compositions and contributes to the understanding of fairy ring ecology.

## Introduction

Soil biodiversity is mostly attributed to its rich contents of microorganisms, but currently most of these microbial taxa remain undescribed, which provides the importance for cultivation and edaphology studies[1]. Sequencing technology is considered as the most effective approach on studying soil microbial diversity, but the sequencing results were differential and usually influenced by different soil processing methods[2]. Moreover, the preservation process of soil samples has been reported to affect microbial DNA extraction, thus directly determining the sequencing quality[3-7]. The internal microbial diversity in soil, as a living biotic environment, after sampling can gradually or drastically change from its original habitat over time. In this case, cryopreservation is one of the common methods applied for soil processing, whereas the impacts of different storage temperature and duration on soil microbial sequencing results were controversial, in which most of the studies supported that only little interference could be introduced[4, 5, 8-11], while few others highlighted the existed interferences on soil microbial sequencing[6, 12]. The divergence in these studies is possibly due to the differences in their research background or the targeted soil types, therefore, all of them possess significance as references on the basis of specific research objectives.

Since the soil storage temperature can influence the quality of high-throughput sequencing[6, 12], there is possibility that cryopreservation for the first few hours after sampling might also have such impact, though extremely low temperature and fast preservation are commonly the best ways to ensure the authenticity of soil samples. As a matter of fact, similar sequencing results were observed previously when comparing the soil samples preserved in liquid nitrogen (<-80°C) and in cooler (−20°C)[11]. However, no study has directly assessed the influence of immediate cryopreservation, which is usually needed for sampling at remote locations. On the other hand, efficient and radical method, with time-saving and high safety properties, in preserving soil samples other than ethanol processing[3, 13], air-drying[8], and freeze-drying[7] is scarce. Drying at high temperature is a promising approach for long-term preservation, which is widespread applied in the biodiversity surveys for microbial DNA of the preserved specimen[14].

The fairy ring is a unique mycological growth pattern usually shown in grasslands and formed by certain fungi living in the soil of special grassland habitat, which has the ecological values in promoting the yield of crops[15]. *Leucocalocybe mongolica*, as one of the representative fairy ring fungi with unique taxonomic status (belongs to monotypic genus), is an incomplete domestication of rare wild mushroom, which is edible, possesses medicinal values, and only lives in fairy soil[16]. Therefore, it is of great significance to investigate fairy ring soil with *L. mongolica* and its microbial compositions, through which we could advance the studies of both fairy rings and *L. mongolica* domestication.

As soil serves as an unstable environment[1, 2], large-scale and multi-point sampling from different geographical environments is crucial for studying the ecological diversity of fairy rings in the soil. Therefore, soil preservation methods, such as high temperature-drying approach, are critical to be evaluated for their influences on detecting the main microbial diversity structures of the soil samples. In addition, the microbial diversity of fairy ring soils could provide important reference values for the subsequent large-scale investigation, due to limited numbers of studies and inconsistent sequencing results[17, 18].

Accordantly, we aimed to discuss the following 3 issues in this study: (i) whether the sequencing results could be affected by immediate *in situ* soil cryopreservation after sampling, (ii) whether the microbial structure in the soil sample could be preserved after high temperature-drying, and (iii) to obtain the microbial diversity information and major species distribution patterns of bacteria and fungi in a fairy ring soil of *L. mongolica*. We used the same fairy ring soil sample and divided it by 3 groups, with 6 biological replicates for each group. The sub-samples were processed with cryopreservation *in situ*, with initial 4 h exposure at room temperature, and with further 12 h drying at high temperature. (S1 Table) 16S rDNA and ITS barcodes for bacteria and fungi analyses and DNA metabarcoding surveys for all the soil samples were further performed. Finally, we discussed the 3 issues mentioned above based on the analyses of microbial community compositions and diversity.

## Materials and methods

### Materials

The soil samples were collected from a *L. mongolica* fairy ring, located in a prairie in Baorixile Town, Hulun Buir City, Inner Mongolia autonomous region, China (49°29′N, 119°59′E, and elevation of 691 m). The pH of sampled soil was 6.4. The sampling area was divided into 3 parts, as Dark zone (Blue flags), Dead zone (Yellow flags), and Outer zone (Red flags), then 8 groups of totally 24 spots were evenly selected for sampling (S1 Fig). For each sampling spot, the soil was collected from the top-layer in a depth of 0-10 cm and a diameter of 5 cm using soil sample collector (S2a Fig). Then, the collected soil sample was screened by using a sieve with 0.3 mm filter (S2b Fig). The soil samples were respectively packed with sterile cryo-storage tubes or paper bags for storage in cold or room temperature conditions, respectively (S2c Fig). The samples were further divided into 3 groups, as DRY (DRY-1 to DRY-6), DI (DI-1 to DI-6), and LN (LN-1 to LN-6), each with 6 biological replicates. Samples in the DRY group were kept at room temperature for 4 h after sampling, followed by flash drying for 12 h at 60°C and long-term storage at room temperature in dry and dark conditions. Samples in the DI group were kept at room temperature for 4 h after sampling, transferred for cryopreservation in dry ice, and finally preserved in cryogenic refrigerator (−80°C) for long-term. Samples in the LN group were immediately transferred into liquid nitrogen for 4 h, then transferred for cryopreservation in dry ice, and finally preserved in cryogenic refrigerator (−80°C) for long-term. (S2d Fig). The timeline of sample processing is shown in Fig 1a.

**Fig 1.**
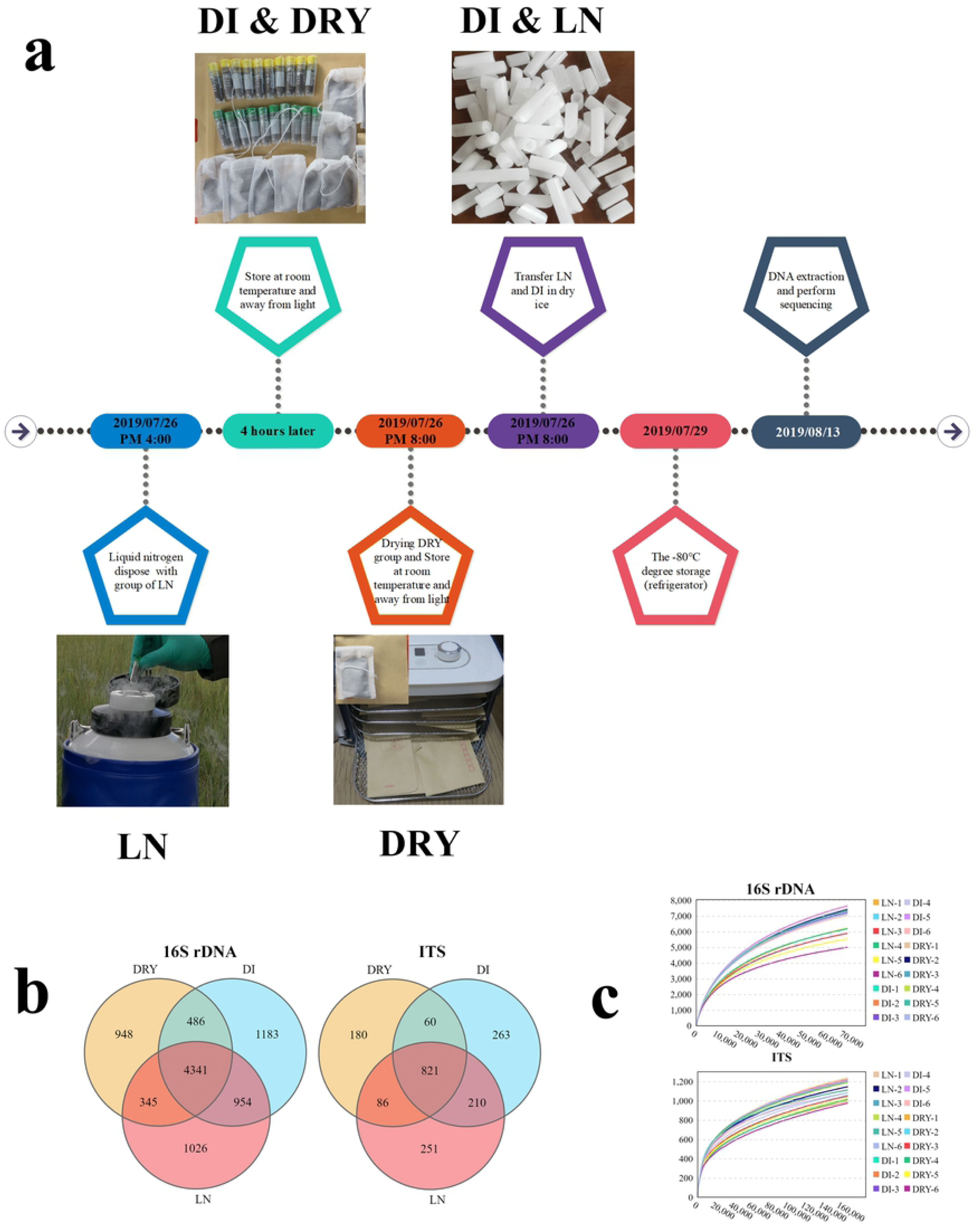
Differences in preservation methods and sequencing results of soil samples across the 3 groups. (a) The timeline of sample processing. (b) The Venn diagram of OTUs from metabarcoding sequencing, in which 16S rDNA represents bacteria and ITS represents fungi. (c) The rarefaction curves of the 18 samples from 16S rDNA and ITS metabarcoding, in which the x-axis represents sequence volume and the y-axis represents OTU numbers.

### DNA extraction, PCR amplification, and sequencing

Microbial DNA was extracted using the HiPure Soil DNA Kits (Magen, Guangzhou, China) according to manufacturer’s protocols. The 16S rDNA V3-V4 region of the ribosomal RNA gene was amplified by PCR, using primers 341F: CCTACGGGNGGCWGCAG; 806R: GGACTACHVGGGTATCTAAT for bacteria[19], and ITS3_KYO2: GATGAAGAACGYAGYRAA; ITS4: TCCTCCGCTTATTGATATGC for fungi[20], following the protocol as follows: 94°C for 2 min, 30 cycles of 98°C for 10s, 62°C for 30s, and 68°C for 30 s, and a final extension at 68°C for 5 min. The PCR reactions were performed in triplicate, in 50 μL mixture containing 5 μL of 10 × KOD Buffer, 5 μL of 2 mM dNTPs, 3 μL of 25 mM MgSO_4_, 1.5 μL of each primer (10 μM), 1 μL of KOD Polymerase, and 100 ng of template DNA. The amplicons were extracted from 2% agarose gels, purified using the AxyPrep DNA Gel Extraction Kit (Axygen Biosciences, Union City, CA, USA) according to the manufacturer’s instructions, and quantified using ABI StepOnePlus Real Time PCR System (Life Technologies, Foster City, CA, USA). The purified amplicons were pooled in equimolar and paired-end sequenced (PE250) on an Illumina platform (Novaseq 6000 sequencing) according to the standard protocols.

### Quality control, reads assembly, and OTUs analysis

(a) Read filtering: Raw data containing adapters or low quality reads were further filtered according to the following rules using FASTP[21]: 1) removing reads containing more than 10% of unknown nucleotides (N) and 2) removing reads containing less than 50% of bases with low quality (Q>value)^20^. (b) Read assembly: The cleaned paired-end reads were merged as raw tags using FLSAH (version 1.2.11)[22] with minimum overlap of 10 bp and mismatch error rate of 2%. (c) Raw tag filtering: Noisy sequences of raw tags were filtered by QIIME (version 1.9.1) pipeline[23] under specific filtering conditions[24] to obtain the high quality clean tags. (d) operational taxonomic units (OTUs) analysis: The effective tags were clustered into of OTU ≥ 97% similarity using UPARSE (version 9.2.64) pipeline[25]. The tag sequence with the highest abundance was selected as the representative sequence within each cluster. Between groups-Venn analysis was performed in R project VennDiagram package (version 1.6.16)[26].

### Community compositional analysis

The representative sequences were classified into organisms by a naive Bayesian model using RDP classifier (version 2.2)[27] based on SILVA database (version 132)[28] for bacterial taxonomy and UNITE database (version 8.0)[29] for fungal taxonomy, with the confidence threshold values ranged from 0.8 to 1. The abundance statistics of each taxonomy were visualized using Krona (version 2.6)[30]. The stacked bar plot of the community composition was visualized in R project ggplot2 package (version 2.2.1)[31]. The heat map of species abundance was plotted using pheatmap package (version 1.0.12, https://cran.r-project.org/web/packages/pheatmap/) in R project (version 2.5.3). The comparison of species among groups was computed by Kruskal-Wallis H test and Welch’s t test in R project Vegan package (version 2.5.3)[32].

### Alpha diversity analysis

Chao1, Simpson, and all other alpha diversity indices were calculated in QIIME (version 1.9.1)[23]. OTU rarefaction curves were plotted in R project ggplot2 package (version 2.2.1)[31]. The comparisons of alpha indices among groups were performed by Kruskal-Wallis H test in R project Vegan package (version 2.5.3)[32].

### Beta diversity analysis

Sequence alignment was performed using Muscle (version 3.8.31)[33], and phylogenetic tree was constructed using FastTree (version 2.1)[34]. Then weighted unifrac distance matrix was generated by GuniFrac package (version 1.0) in R project[35]. The NMDS of weighted unifrac distances was generated in R project Vegan package (version 2.5.3)[32] and plotted in R project ggplot2 package (version 2.2.1)[31]. The statistical analyses of Kruskal-Wallis H test and Anosim test were calculated in R project Vegan package (version 2.5.3)[32].

### Data Availability

The raw amplicon sequencing dataset is available in the NCBI Sequence Read Archive under BioProject ID:PRJNA635768.

## Results

### Sequencing Analysis

A total of 5,350,348 metabarcoding tags were obtained from sequencing data. The clustering analyses of the soil samples showed that there were 7171 and 1161 OUTs in average for bacteria (based on 16S rDNA) and fungi (based on ITS), respectively (Table 1). The Venn diagram exhibited that 46.76% (4341/9283) bacterial and 43.88% (821/1871) fungal OTUs were shared among the 3 groups of soil samples (Fig 1b). The leveled-off rarefaction curves of all the soil samples for both bacteria and fungi indicated the near saturation of the metabarcoding sequencing (Fig 1c). Generally, we found that the OTUs number of bacteria in the fairy ring soil was significantly higher than that of fungi.

**Table 1.** General metabarcoding sequencing results of the soil samples.

| Sample ID | 16S rDNA |  |  | ITS |  |  |
| --- | --- | --- | --- | --- | --- | --- |
|  | Tags | N90 (bp) | OTUs | Tags | N90 (bp) | OTUs |
| LN-1 | 105391 | 441 | 7721 | 183533 | 352 | 1271 |
| LN-2 | 103243 | 441 | 7528 | 185974 | 351 | 1181 |
| LN-3 | 94364 | 441 | 7417 | 179043 | 349 | 1221 |
| LN-4 | 96054 | 441 | 7297 | 178074 | 351 | 1211 |
| LN-5 | 98194 | 441 | 7456 | 200309 | 350 | 1161 |
| LN-6 | 108598 | 441 | 7744 | 172559 | 351 | 1091 |
| DI-1 | 99046 | 441 | 7992 | 248485 | 352 | 1325 |
| DI-2 | 104394 | 441 | 7927 | 189108 | 352 | 1265 |
| **DI-3** | 102727 | 441 | 7942 | 193608 | 352 | 1193 |
| **DI-4** | 108138 | 441 | 7705 | 188333 | 349 | 1123 |
| **DI-5** | 99954 | 441 | 7723 | 168489 | 351 | 1212 |
| **DI-6** | 104372 | 441 | 7452 | 203033 | 351 | 1185 |
| **DRY-1** | 104807 | 441 | 6654 | 243722 | 340 | 1121 |
| **DRY-2** | 106470 | 441 | 6352 | 210081 | 340 | 1144 |
| **DRY-3** | 109328 | 441 | 6357 | 191669 | 342 | 1094 |
| **DRY-4** | 99839 | 441 | 6501 | 187538 | 344 | 1054 |
| **DRY-5** | 107343 | 441 | 5932 | 183210 | 340 | 1052 |
| **DRY-6** | 107876 | 441 | 5386 | 183442 | 340 | 1009 |
| **MEAN** | 103341 | 441 | 7171 | 193900 | 347 | 1161 |

### Microbial community composition

Based on SILVA[28] and UNITE[29] databases, we analyzed the differences in the community compositions of soil bacteria and fungi among LN, DI, and DRY groups. Fig 2 shows the microbial taxonomic structures for soil samples at the levels of phylum and family. For bacteria in LN, DI, and DRY groups (Fig 2ac, S2 and S3 Tables), the top 10 most abundant phyla were Actinobacteria (22.3, 24.72, and 24.25%), Verrucomicrobia (22.91, 16.69, and 19.22%), Proteobacteria (15.93, 18.64, and 19.64%), Planctomycetes (16.2, 14.59, and 13.75%), Acidobacteria (10.25, 11.79, and 11.17%), Chloroflexi (3.74, 4.07, and 3.51%), Gemmatimonadetes (3.27, 3.63, and 3.73%), Bacteroidetes (1.43, 1.71, and 1.6%), Rokubacteria (0.65, 0.74, and 0.8%), and Patescibacteria (0.91, 0.86, and 0.4%); while the top 10 most abundant families included Chthoniobacteraceae (20.68, 15.02, and 17.78%), Gemmataceae (6.54, 5.55, and 6.52%), Pyrinomonadaceae (3.5, 4.09, and 3.52%), Tepidisphaerales sp. (3.91, 3.66, and 3.0%), Gemmatimonadaceae (2.95, 3.29, and 3.49%), Xanthobacteraceae (2.66, 2.92, and 3.82%), Sphingomonadaceae (2.34, 2.98, and 3.57%), Solirubrobacteraceae (2.32, 2.45, and 2.48%), Solirubrobacterales sp. (2.22, 2.45, and 2.54%), and Pirellulaceae (2.92, 2.35, and 1.89%). For fungal compositions in LN, DI, and DRY groups (Fig 2bd, S4 and S5 Tables), Basidiomycota (81, 82.1, and 68.49%), Ascomycota (17.04, 16.3, 29.04%), and Anthophyta (1.2, 1.04, and 0.97%) were the dominant phyla; while Tricholomataceae (76.99, 78.41, and 65.05%), Geoglossaceae (1.11, 1, and 1.42%), Aspergillaceae (0.92, 0.79, and 1.45%), Clavariaceae (1.1, 1.02, and 0.76%), and Helotiales sp. (0.57, 0.56, and 1.02%) were the dominant families. Overall, we detected similar microbial compositional distribution patterns among the 3 groups of fairy ring soil.

**Fig 2.**
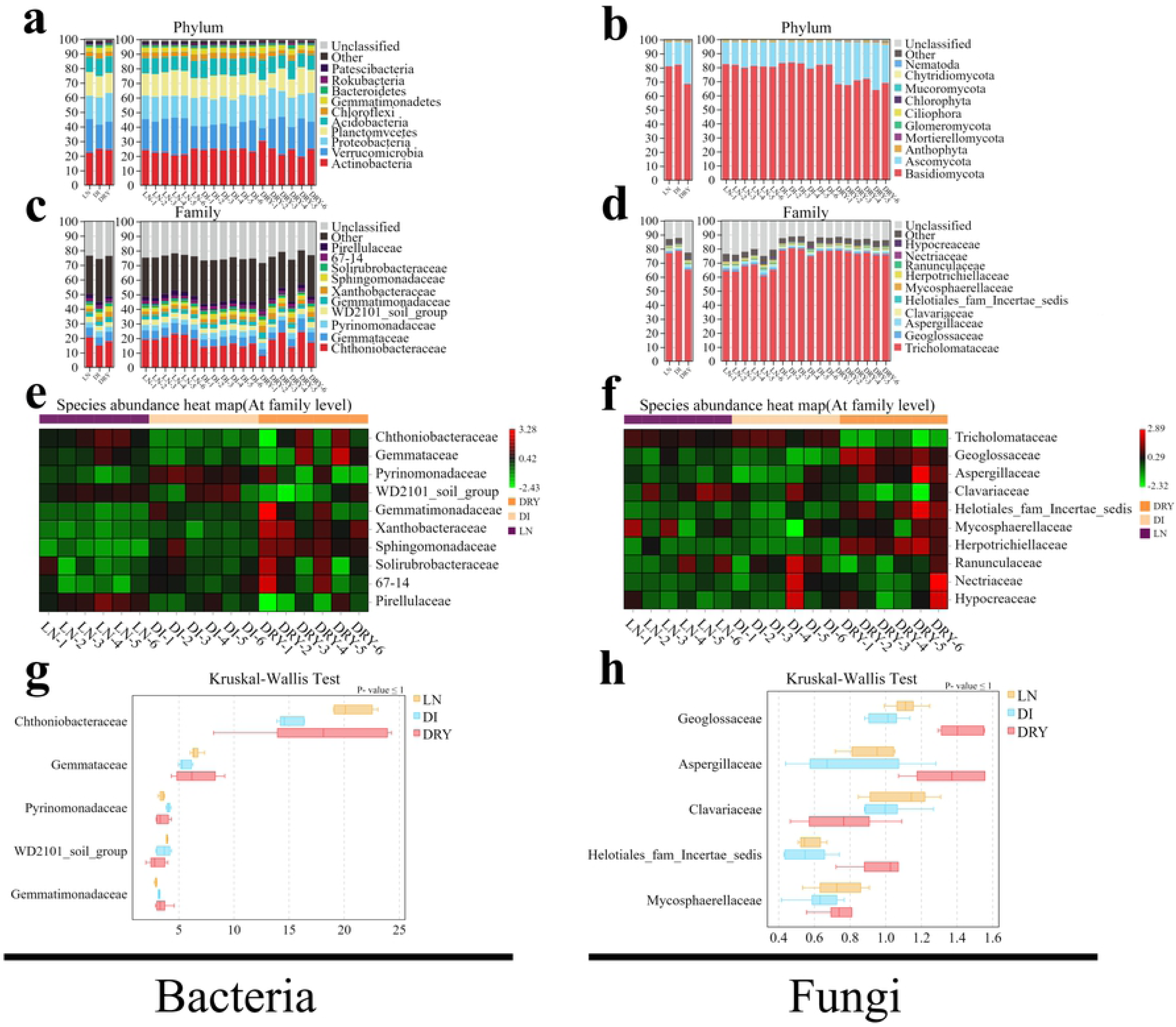
OTUs taxonomic analysis of the soil samples across the 3 groups. (a-d) Taxonomy of bacteria (a,c) and fungi (b,d) at phylum (a,b) and family (c,d) levels, in which the x-axis represents the sample ID and the y-axis represents the relative abundance. (e,f) Heat map of the top 10 families of bacteria (e) and fungi (f), in which the x-axis represents the sample ID and the y-axis represents the family. (g,h) The Kruskal-Wallis test of the top 5 families of bacteria (g) and fungi (h), in which the x-axis represents the relative abundance and the y-axis represents the family. Tricholomataceae was excluded from the results of fungi.

However, based on the heat maps (Fig 2ef), when species were compared separately across all soil samples, we observed differences in the microbial abundance among individual samples as well as across groups. The relative abundances of the top 10 families of both bacteria and fungi in the DRY group exhibited various distribution patterns, which indicates that high temperature-drying process might affect the constancy of soil microbial sequencing. On the other hand, the relative abundances in 3 of the top 5 families, i.e. Chthoniobacteraceae, Gemmataceae, and Pirellulaceae, in the LN group were higher than those in the DI group, whereas higher abundance of Pyrinomonadaceae was found in the DI group (Fig 2e), indicating that the difference in 4 h initial preservation of soil samples could influence their microbial sequencing consistency. Moreover, the relative abundance of Tricholomataceae was lower in the DRY group when compared with those in LN and DI groups, while 4 of the top 5 families, i.e. Geoglossaceae, Aspergillaceae, and Helotiales sp., were relatively higher in the DRY group but Clavariaceae showed the opposite result.

Additionally, in order to further clarify the differences we observed by visualizaion in the heat map, the Kruskal-Wallis and Welch’s t-tests of bacteria (Fig 2g) and fungi (Fig 2h) were performed. The Kruskal-Wallis test result revealed that most of the top abundant families were insignificantly different and two of the dominant families of bacteria and fungi, i.e. Chthoniobacteraceae and Tricholomataceae, showed the biggest difference, in which Chthoniobacteraceae (P-value = 0.038) exhibited 5.66% differences in DI (LN:DI) and 2.9% differences in DRY (LN:DRY), while Tricholomataceae (P-value = 0.002) exhibited −1.43% differences in DI (LN:DI) and 11.94% in DRY (LN:DRY). The Welch’s t-test revealed that, when comparing the difference between LN and DI groups, 3 of the top 5 bacterial families exhibited significant difference, including Gemmataceae (0.99% difference, P-value = 0.008), Pyrinomonadaceae(−0.59% difference, P-value = 0.001), and Gemmatimonadaceae(−0.54% difference, P-value = 0.004) (S3a Fig); when comparing difference between LN and DRY groups, 4 of the top 5 fungal families exhibited significant difference, including Geoglossaceae (−0.31% difference, P-value = 0.001), Aspergillaceae (−0.53% difference, P-value = 0.02), Clavariaceae (0.34% difference, P-value = 0.02), and Helotiales sp. (−0.45% difference, P-value = 0.003) (S3b Fig). To be noted, the fungal family Tricholomataceae was excluded from the Kruskal-Wallis test due to its exorbitant abundance. Besides, soil samples in the DRY group were associated with less repeatability in bacterial abundances (such DRY-1,2 vs DRY-3,5; Fig 2e), and less differences in fungal abundances were observed between LN and DI groups (Fig 2f). Therefore, the Welch’s t-tests for bacterial abundance of LN vs DRY and for fungal abundance of LN vs DI were not performed. Overall, the abundance pattern of fungal community was moderately affected by different soil preservation methods, but the pattern of soil bacterial community was significantly influenced by different preservation methods.

### Alpha diversity analysis

We further sought to better understand the changes of both bacterial and fungal diversity in the soil samples introduced by *in situ* cryopreservation. As shown in Fig 3 and Table 2, a total of 6 alpha diversity indices were estimated, including Sobs (observed species, equal to the number of OTUs), Chao1, Ace, Shannon, Simpson and Goods_coverage. For bacteria, among all 18 soil samples, we detected Sobs values at a range of 5386 (DRY-6) to 7992 (DI-1), Chao1 from 7204 (DRY-6) to 10530 (DI-1), Ace from 6857 (DRY-6) to 10537 (DI-3), Shannon from 9.92 (DRY-5) to 10.72 (DI-1), and Simpson and Goods_coverage indices all exceeded 0.95 (Table 2). Among the 6 indices, Sobs (P-value = 0.002), Chao1 (P-value = 0.002), and Ace (P-value = 0.002) were significantly different across soil groups, in which the DRY group exhibited the lowest values (Fig 3a and Table 2), suggesting that drying soil at high temperature could reduce both the richness and diversity of bacterial species. For fungal alpha diversity, we observed Sobs ranging from 1009 (DRY-6) to 1325 (DI-3), Chao1 from 1450 (DRY-6) to 1822 (DI-1), Ace from 1463 (LN-6) to 1771 (LN-1), Shannon from 2.48 (DI-2, DI-3) to 3.83 (DRY-5), Simpson from 0.4 (DI-2, DI-3) to 0.65 (DRY-5), and Goods_coverage values all exceeded 0.99 (Table 2). The soil samples in the DRY group also had the lowest Sobs value (P-value = 0.01) but the highest Shannon value (P-value = 0.002), indicating that the high temperature-drying process could reduce the richness of fungal species but increase their diversity (Fig 3b and Table 2).

**Table 2.**
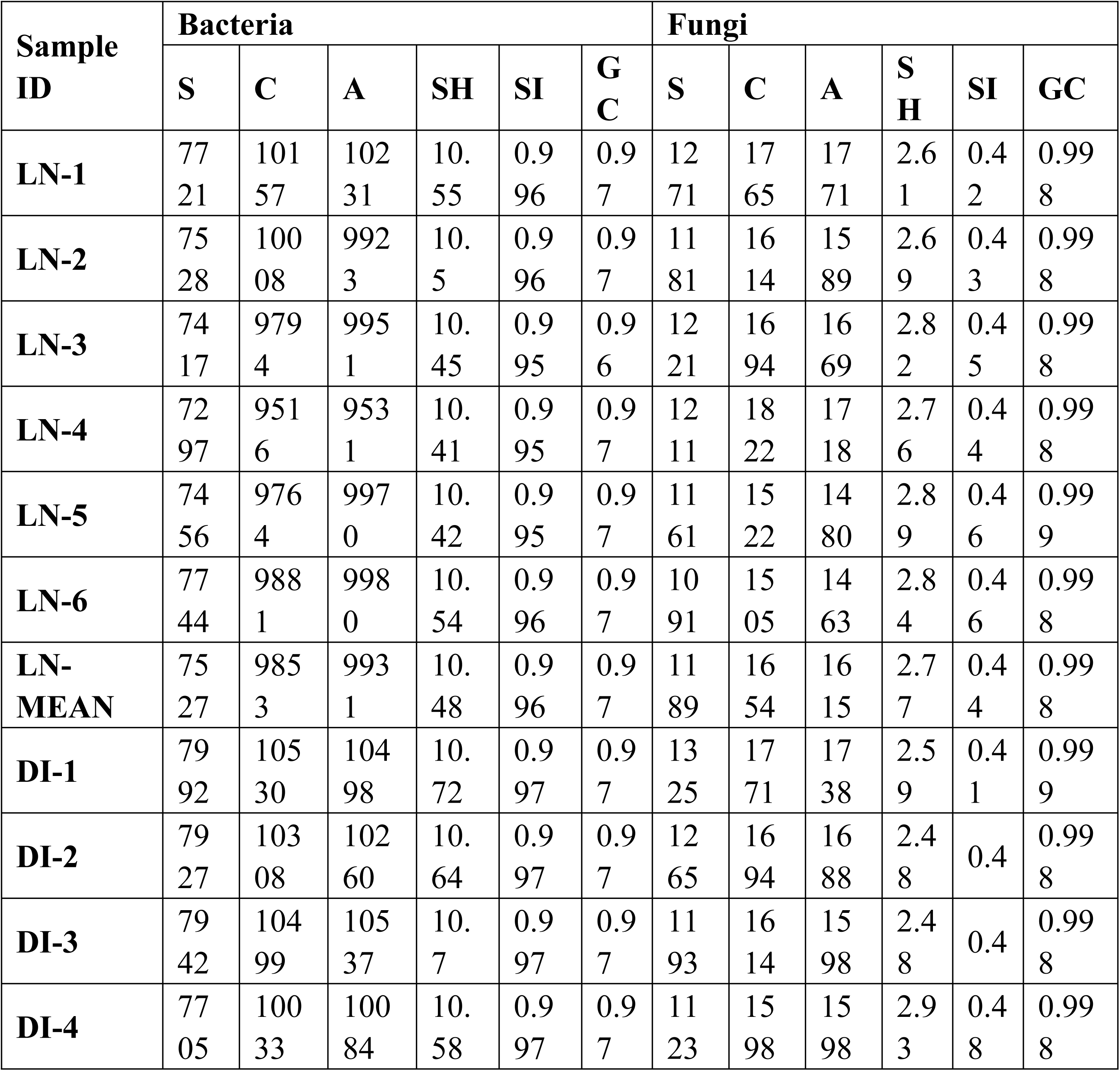
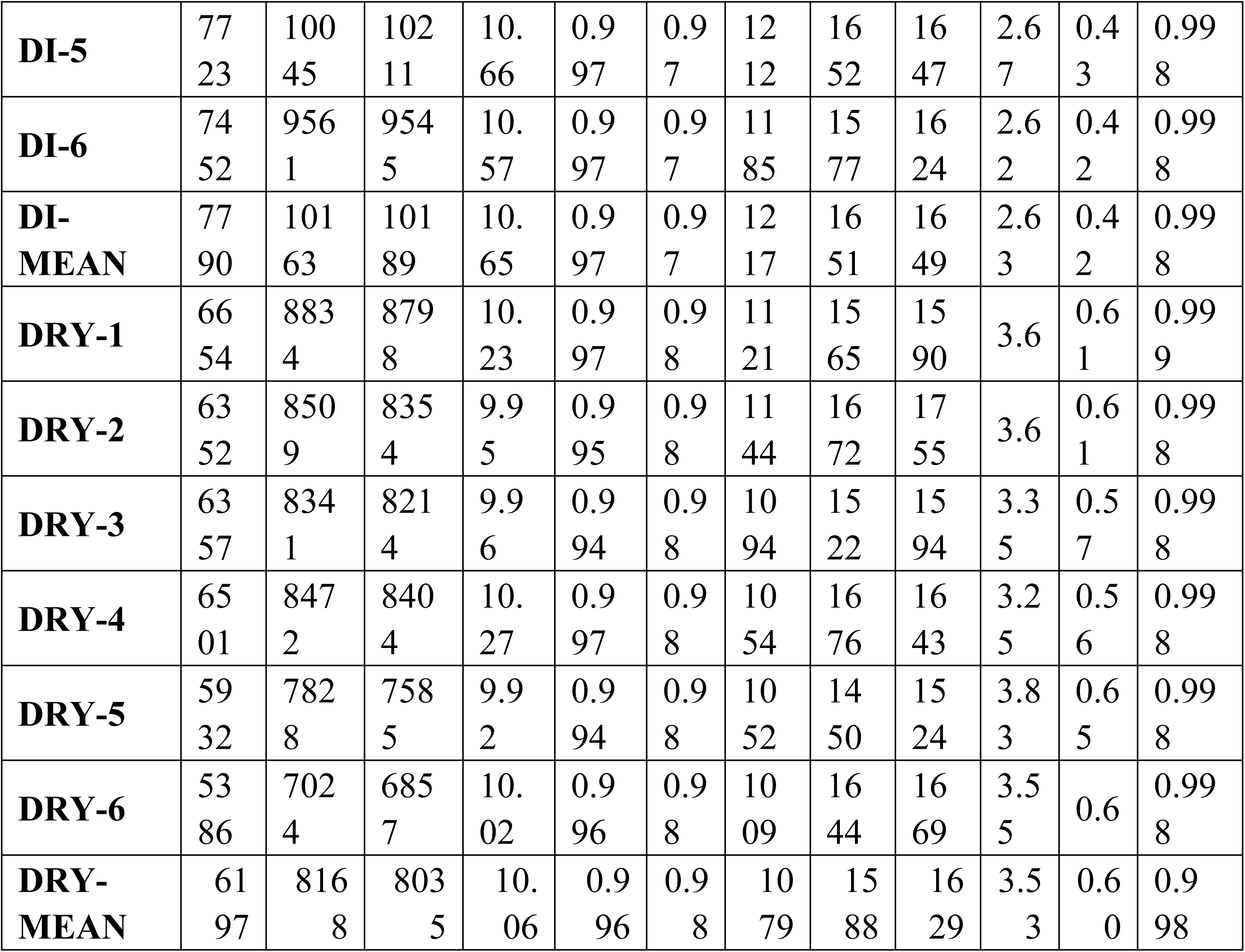
Alpha diversity indices for the soil samples. Note: S = Sobs, C = Chao1, A = Ace, SH = Shannon, SI = Simpson, GC = Good’s Coverage.

**Fig 3.**
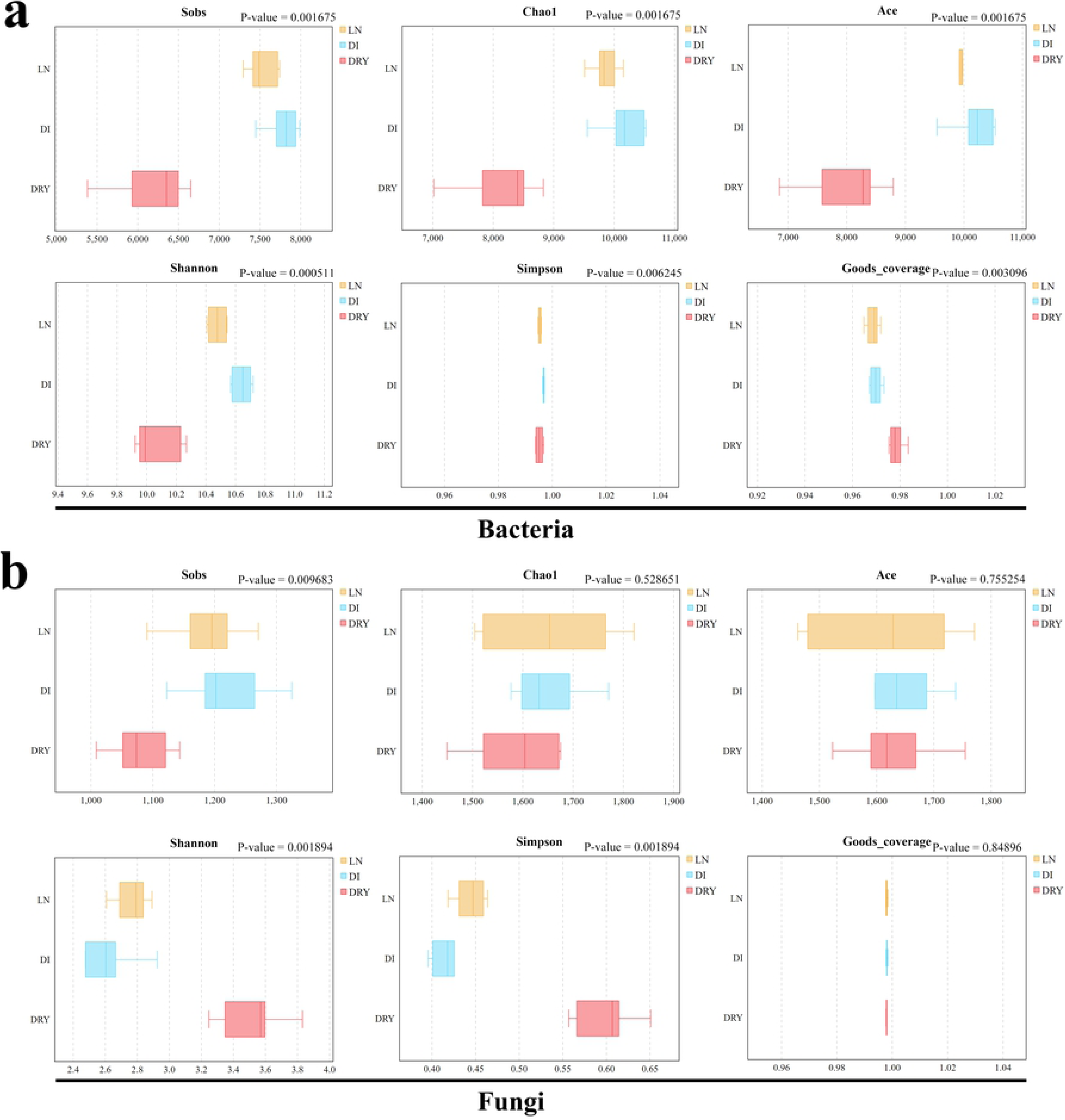
Inter-group analysis of alpha diversity. The x-axis represents the indices of alpha diversity and the y-axis represents the groups of soil samples.

### Beta diversity analysis

Finally, beta diversity analysis was performed in order to understand the variation of soil microbes within individual group and across different groups. The beta diversity based on weighted unifrac distance index from OTUs is shown in Fig 4. For bacteria, as shown in the heat map (Fig 4a), the soil samples in LN and DI groups clustered in their own branches, while the samples in the DRY group failed to cluster, and these findings were further confirmed by the non-metric multi-dimensional scaling (NMDS) analysis shown in Fig 4c. The analysis of similarities (Anosim), shown in Fig 4e, also revealed that, compared with the DRY group, LN and DI soil groups had lower intra-group differences (R-value = 0.573, P-value < 0.001), inferring the moderate inter- and intra-group differences across the 3 groups,. For fungi, as Fig 4d shown, we observed that the soil samples clustered into 2 branches, in which the LN group clustered with the DI group, while the DRY group clustered alone (Fig 4b). Similar results were found in the NMDS analysis (Fig 4d). The P-value of Anosim was less than 0.001 (Fig 4f), which indicates that the diversity differences in soil fungi among groups were greater than those in soil bacteria. Overall, soil bacterial diversity in LN and DI groups formed 2 complete clusters, and compared with the DRY group, they contained less intra-group difference; for fungi, soil bacterial diversity in LN and DI groups could not be significantly distinguished by OTU genetic distance, but they were clustered separately from the DRY group.

**Fig 4.**
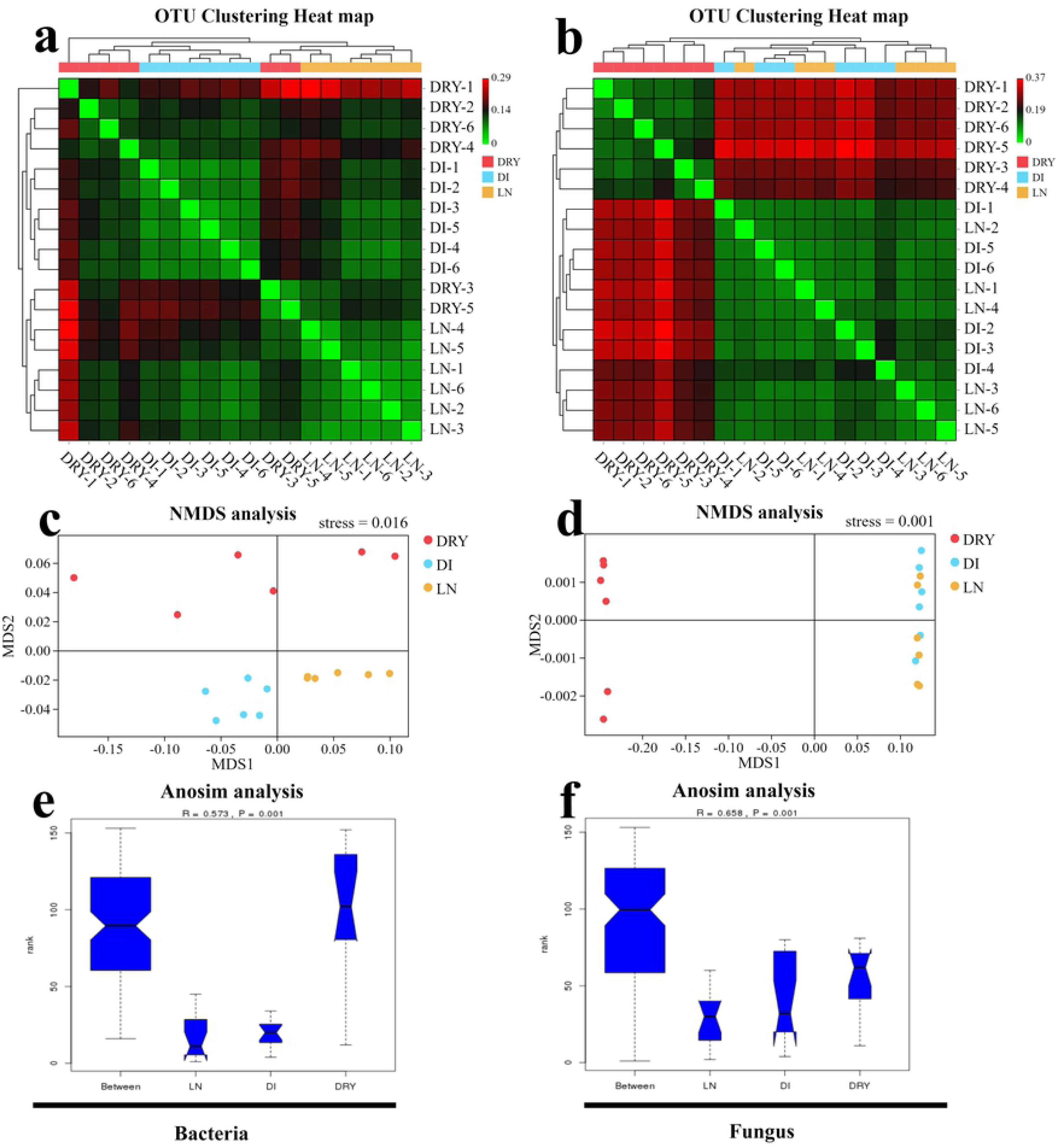
Beta diversity analysis of the soil samples across the 3 groups. (a,b) Heat map of weighted unifrac distance index based on UPGMA clustering from bacteria (a) and fungi (b), in which the bottom x-axis and right y-axis represent the sample ID, and the upper x-axis and left y-axis represent UPGMA clustering results. (c,d) The NMDS plot of bacteria (c) and fungi (d), in which MDS1 represents abundance distance matrix and MDS2 represents evolutionary distance matrix. (e,f) The map of Anosim analysis, in which the x-axis represents the group ID, the y-axis represents the relative difference grade, the R value represents the difference level, and the P value represents the significance of the difference.

## Discussion

In this study, we performed community compositional analysis as well as alpha and beta diversity analyses to assess the influences of 3 different preservation processes of soil samples on their microbial DNA metabarcoding sequencing results. We found that the initial 4 hours of soil processing were important for the quality of microbial sequencing, whereas high temperature-drying could not seriously influence the sequencing results. Meanwhile, we have also obtained abundant background information on soil microbial composition and diversity in the habitat environment of the fairy ring of *L. mongolica*.

Four-hour soil processing at room temperature could influence the microbial sequencing. First, we observed that the soil microbial community of the DI group was different compared with the LN group, in terms of the abundance of major distributed families, especially Chthoniobacteraceae, Gemmataceae, Solirubrobacterales sp., and Pirellulaceae, and differences were also found within the biological replicates (Fig 2eg). Secondly, we detected that the soil bacteria in the DI group diverged significantly from those in the LN group through UPGMA clustering analysis (Fig 4a), with all biological replicates from both groups clustered into their own branches. Finally, the analyses of NMDS (Fig 4c) and Anosim (Fig 4e) validated the differences between LN and DI groups. Previously, Schnecker found that when soil samples were stored at 4°C, the biomarker bacterial populations were changed[12], and similarly we also revealed that if the soil samples were not immediately *in situ* cryo-preserved, the sequencing results of soil bacterial diversity could be changed, by as short as 4 hours initial storage at room temperature. However, for fungi, we did not find any substantial influence of initial room temperature-preservation on the quality of DNA metabarcoding sequencing, indicated by the similar community fungal compositions and diversity in LN and DI groups (Fig 2bdfh, 3b, and 4bd), which is consistent with the studies by Cui[5]. In addition, the Ace index in bacteria showed that the LN group contained less intra-group difference within the biological replications than that of the DI group (Fig 3a). Since the Ace index could cover the microbial species with low abundance (OTUs number below 10 but more than 1)[23], this result indicates that the low abundance-species in the soil samples were well preserved by *in situ* cryopreservation. Furthermore, our study is related to a recent study by Delavaux[11], which also supported that low temperature-preservation of soil samples is important for the accuracy of microbial metabarcoding sequencing. However, compared with the study by Delavaux, our study focused more on the influence of the temperature during initial soil sample processing instead of the difference introduced by low temperature-storage (cooler or liquid nitrogen). In general, the community composition in the soil samples was influenced by the initial preservation method, but at a low degree (Fig2. ab), possibly due to the shared soil source in LN and DI groups. At the same time, given that the initial duration of sample processing in the DI group was very short, the microbial differences between LN and DI groups also revealed the accuracy of our sequencing results.

High temperature-drying could not influence the major microbial community. We have observed that the major microbial community in the soil samples was not significantly changed by 12 hours drying at 60°C, reflected by the shared similar relative abundances of the top 5 bacterial and fungal families between the DRY soil group and LN/DI soil group (Fig 2gh). As a matter of fact, previous studies have investigated the influences of long time storage, normal drying, and freeze-drying on soil microbial community[3, 5, 7-10], and here our finding demonstrates that high temperature-drying can serve as an efficient approach for soil sample processing, although the drying process reduced microbial alpha diversity (Fig 3) and leads to less repeatability in the analyses of soil bacteria and fungi (Fig 2ef and 4ae). Overall, the approach of high temperature-drying is only limited in preserving the major microbial community composition of the soil samples, whereas the method could not ensure high-precision of the sequencing results. In addition, this approach is also not suitable for ensuring high repeatability within biological replicates and preserving microbial species with low abundances. Therefore, *in situ* cryopreservation is a more ideal processing method for high-throughput sequencing study of soil microbial ecosystem.

The basic information of microbial composition and diversity in soil of fairy ring ecosystem. Through DNA metabarcoding sequencing analysis with the barcodes of 16S rDNA and ITS in this study, we obtained a basic background information of the soil microorganisms in fairy ring of *L. mongolica* and identified the major distribution patterns of both bacteria and fungi in the soil. First, we observed that the total number of OTUs obtained from the soil samples was close to the maximum number of Chao1 and Ace indices for both bacteria and fungi, indicating that a nearly complete DNA barcoding background information of soil microbes was obtained through 18 repeated sequencing (Fig 1b and Table 2). In related to a relevant studies, we were able to obtain higher numbers of bacterial OTUs (ranging from 5386 to 7992; Table 1) in fairy ring of *L. mongolica* compared with 348 to 569 OTUs sequenced from the fairy ring soil of *Floccularia luteovirens*[18], and we showed a similar fungal OTUs in fairy ring of *L. mongolica* (ranging from 1009 to 1325; Table 1) with the OTUs in the fairy ring soil of *Agaricus gennadii* (ranging from 1242 to 1398)[17]. These suggest that the OTU numbers of soil bacteria were more unstable than those of soil fungi. Second, we found that the bacteria and fungi in the fairy ring soil were dominated by the family of Chthoniobacteraceae and Tricholomataceae, respectively, whose abundances were much higher than the following families (Gemmataceae and Geoglossaceae, respectively) in all the soil samples. In fact, Chthoniobacteraceae in the phylum of Verrucomicrobia was previously reported to be one of the dominant soil bacterial taxa based on 16S rRNA studies and its relative abundance in average was between 0% and 20% in various soil environments[36, 37]. Moreover, Chthoniobacteraceae is also one of the major bacterial families in several other environments, such as mine area and water[38, 39], and it contributed in nitrogen metabolism of certain ecosystems[40], potentially indicating its important functions in fairy ring ecology. On the other hand, Tricholomataceae exhibited extremely high abundance in all the soil samples in this study (Fig 2b), from which we speculated that the dominance was attributed to the mycelium of *L. mongolica* during its fruiting body formation.

## Conclusion

In this study, we assessed the importance of *in situ* cryopreservation and high temperature-drying methods for fairy ring soil sample processing. Then we analyzed the microbial structure for both the distribution pattern and relative abundance of the soil samples from *L. mongolica* fairy ring, which could provide references for further ecological studies on fairy ring ecosystem and *ex situ* conservation of *L. mongolica*.

## Supporting information

**S1 Fig. The soil sampling spots in the fairy ring of *Leucocalocybe mongolica***. The flag of blue, yellow, and red represent sampling spots of Dark zone, Dead zone, and Outer zone, respectively.

**S2 Fig. The soil sampling process**. (a) The sampling size and method. (b) The screening method. (c) The packing method. (d) The soil sample processing in the LN group.

**S3 Fig. Welch’s t test analysis in top 7 abundant families in bacteria and fungi**. Subfigure a represents families of bacteria and b represents families of fungi. The left half of the figure shows the ordinate of different species. X-axis shows the mean abundance of species. The difference value of the abundance between groups is shown in the right half of the x-axis. The point color represents the group with higher abundance, the error bar of the point represents the fluctuation range of the 95% confidence interval of the difference value, and y-axis represents the significance (p-value) of the difference between groups of the corresponding families.

**S1 Table. Differences in sampling methods among the three groups**.

**S2 Table. The phylum level bacterial OTU classification**. The values represent the number of taxonomic tags.

**S3 Table. The family level bacterial OTU classification**. The values represent the number of taxonomic tags.

**S4 Table. The phylum level fungal OTU classification**. The values represent the number of taxonomic tags.

**S5 Table. The family level fungal OTU classification**. The values represent the number of taxonomic tags.

## Acknowledgements

This study was financially supported by The Biodiversity Survey and Assessment Project of the Ministry of Ecology and Environment, China (No.2019HJ2096001006). We acknowledge TopEdit (www.topeditsci.com) for the linguistic editing and proofreading during the preparation of this manuscript.

## Author Contributions

BT conceived of the study. MZD performed research and wrote the manuscript.

